# Adolescent intermittent ethanol exposure effects on kappa opioid receptor mediated dopamine transmission: Sex and age of exposure matter

**DOI:** 10.1101/2020.06.17.157925

**Authors:** Mary B. Spodnick, Raymond T. Amirault, Trevor T. Towner, Elena I. Varlinskaya, Linda P. Spear, Anushree N. Karkhanis

## Abstract

Underage alcohol drinking increases the risk of developing alcohol use disorder (AUD). In rodents, adolescent ethanol exposure augments ethanol consumption and anxiety-like behavior while reducing social interaction. However, the underlying mechanisms driving these adaptations are not understood. The dopamine and kappa opioid receptor (KOR) systems in the nucleus accumbens (NAc) are implicated in affective disorders and AUD, with studies showing augmented KOR function and reduced dopamine transmission in ethanol-dependent adult animals. Thus, this study, we examined the impact of adolescent intermittent ethanol (AIE) exposure on dopamine transmission and KOR function. Rats were exposed to water or ethanol (4 g/kg, intragastrically) every-other-day during early (PD25–45) or late (PD45–65) adolescence. While AIE exposure during early-mid adolescence (early AIE) did not alter dopamine release in male and female rats, AIE exposure during late adolescence (late AIE) resulted in greater dopamine release in males and lower dopamine release in females. To determine the impact of AIE exposure on KOR function, we bath applied cumulative concentrations of KOR agonist, U50,488 (0.01–1.0 μM), and measured its effect on dopamine release. Early AIE exposure potentiated KOR-mediated inhibition of dopamine release in female rats, while late AIE exposure attenuated this effect in male rats. Together these data suggest that AIE-exposure impact on neural processes is dependent on sex and exposure timing. These differences likely arise from differential developmental timing in males and females. This is the first study to show changes in KOR function following AIE exposure.

## INTRODUCTION

Adolescence is a key period for initiation of alcohol use [1] and the beginning of the development of alcohol use disorder (AUD) [2]. Approximately 58% of teenagers report having at least one alcoholic drink prior to reaching 18 years of age [3]. Moreover, about 4.7 % of adolescents (ages 12 – 17) and about 35% of young adults (ages 18 – 25) engaged in binge drinking in the past month [3]. Initiation of binge drinking patterns before the age of 14 is *especially* concerning given that the prevalence of alcohol dependence following early initiation (before 14 years) is four times greater than if individuals wait until the legal age limit [4–7]. Earlier age of initiation of alcohol consumption is also correlated with mental health issues, including AUD, later in life [2,8,9].

Similar to humans, adolescent ethanol exposure in rodents promotes long-lasting maladaptive behaviors [10], including augmented anxiety-like behavior [11,12] and attenuation of social investigation and preference [13]. These changes are differentially affected by age of exposure in males and females; specifically, while both early adolescent (PD 25 – 45) and late adolescent (PD 45 – 65) exposures result in greater anxiety-like behavior in males, only early exposure females exhibit augmented anxiety-like behavior [12]. With respect to social interaction, social investigation and social preference were significantly reduced in early-exposed males, with no social alterations evident in late-exposed males and females regardless of exposure timing [13]. Social interaction has a substantial reward value, with mesolimbic (ventral tegmental area [VTA] to nucleus accumbens [NAc]) dopamine projections being directly involved in modulation of social reward [14,15]. Thus, it is possible that the adolescent alcohol exposure-associated social deficits and enhanced anxiety-like behaviors may be driven by neurochemical changes in the NAc, particularly since early-mid adolescence is a critical time for maturation of mesolimbic dopamine neurons [16,17].

Rodent studies have demonstrated a direct relationship between ethanol and dopamine, with acute ethanol challenge resulting in activation of dopamine neurons in the VTA [18,19] subsequently elevating extracellular dopamine levels in the NAc core of adult male rats [20]. In contrast, chronic intermittent ethanol (CIE) exposure in adult male mice attenuates stimulated dopamine release during acute withdrawal in the NAc core [21,22]. These CIE-mediated reductions in dopamine have been shown to last for up to 72 hours after the cessation of final exposure [21–23]. Relatively fewer studies, however, have explored ethanol’s impact on dopamine transmission following adolescent exposure. There is some evidence that adolescent ethanol exposure between postnatal days (PD) 30 and 45 can decrease accumbal dopamine tissue content in male mice when assessed following five days of abstinence [24]. Interestingly, intermittent ethanol exposure between PD 30 and 50 followed by 15 days of abstinence resulted in elevated basal levels of dopamine in the NAc in adult rats [25]. On the other hand, a protracted abstinence of 25 days following intragastric intermittent ethanol exposure during early adolescence (PD 25 – 45) revealed no differences in stimulated dopamine release at baseline in NAc core of adult rats [26]. Although there is some overlap in age of exposure across these studies, period of abstinence and the 5-day shift in exposure time may drive the differences observed as maturation of the dopamine system is at its peak during this time [16,17]. The focus of this study, however, is to systematically compare the impact of ethanol exposure during early-mid adolescence and late adolescence/emerging adulthood [27,28]. These age spans correspond to approximately 10–18 and 18–25 years of age in humans, respectively [28].

Among the various mechanisms that regulate dopamine release in the NAc [29], the inhibitory control of KORs is of particular importance given that it is also a target of ethanol. Specifically, KORs are located on the terminals of accumbal dopamine neurons [30] and activation of these receptors inhibits dopamine release [21,22,31–33]. This KOR-mediated inhibition of dopamine release in the NAc core is enhanced in alcohol dependent adult male mice [21,22], and systemic KOR blockade, in turn, reverses dependence-induced enhancement in ethanol consumption in mice [21] and rats [34]. Conversely, adult male rats with moderate drinking history show increased ethanol intake in response to KOR inhibition [35]. Adding to the complexity of this picture is that adult female rats with moderate drinking history show reduced ethanol intake following KOR blockade, whereas KOR manipulation does not affect ethanol intake in adolescent male and female rats [35]. Given the relationship between KORs and dopamine illustrated by these alcohol exposure models, it is possible that adolescent intermittent ethanol (AIE) exposure may lead to sex- and age-dependent disruptions in dopamine transmission via changes in KOR function.

While there is paucity in literature examining AIE-mediated neurobiological changes, studies measuring affect-related behavioral effects show that AIE-induced behavioral adaptations were dependent on time of exposure, either early adolescence (PD 25 – 45) or late adolescence/emerging adulthood (PD 45 – 65), and sex [12,13]. Thus, in this study we examined the impact of AIE exposure during early and late adolescence followed by forced protracted abstinence in male and female rats on dopamine transmission and KOR function using *ex vivo* fast scan cyclic voltammetry (FSCV). Based on previous behavioral and neurobiological data, we hypothesized that AIE exposure during early adolescence will not alter dopamine efflux at baseline; however, late adolescent exposure will only attenuate dopamine release in males. Moreover, we predicted that KOR function will be potentiated following AIE exposure during early adolescence in both sexes, but only in males exposed to AIE during late adolescence. This is the first study to examine the effects of AIE exposure on KOR function in the NAc.

## METHODS

### Experimental Subjects

Male and female Sprague-Dawley rats derived from our breeding colony at Binghamton University were used. On PD 1, litters were preferentially culled to a ratio of 6 males and 4 females when possible. On PD21, rats were weaned and housed in pairs with a same-sex littermate. In order to limit the possible confound of litter effect [36], one rat per sex per litter was assigned to each experimental group. Rats were maintained on a 12:12-h light/dark schedule (lights on at 700 hours) with standard rat chow (LabDiet 5L0D – PicoLab Laboratory Rodent Diet, ScottPharma Solutions, Marlborough, MA) and tap water available ad libitum. Maintenance and experimental treatment of rats was in accordance with the National Institutes of Health guidelines for animal care using procedures approved by the Binghamton University Institutional Animal Care and Use Committee.

### Adolescent intermittent ethanol exposure

Rats were exposed to ethanol (4 g/kg, 25% v/v) via intragastric gavage (i.g.) during early/mid (early AIE, PD 25 – 45) or late (late AIE, PD 45 – 65) adolescence. Age- and sex-matched controls were administered an isovolumetric amount of water. Animals were given ethanol or water every other day during the light phase between 1300 – 1600. An intermittent pattern of exposure was used to better represent human alcohol consumption [37]. Experimental timelines can be seen in **Figures 1A** (early-AIE exposure) and **1B** (late-AIE exposure).

**Figure 1:**
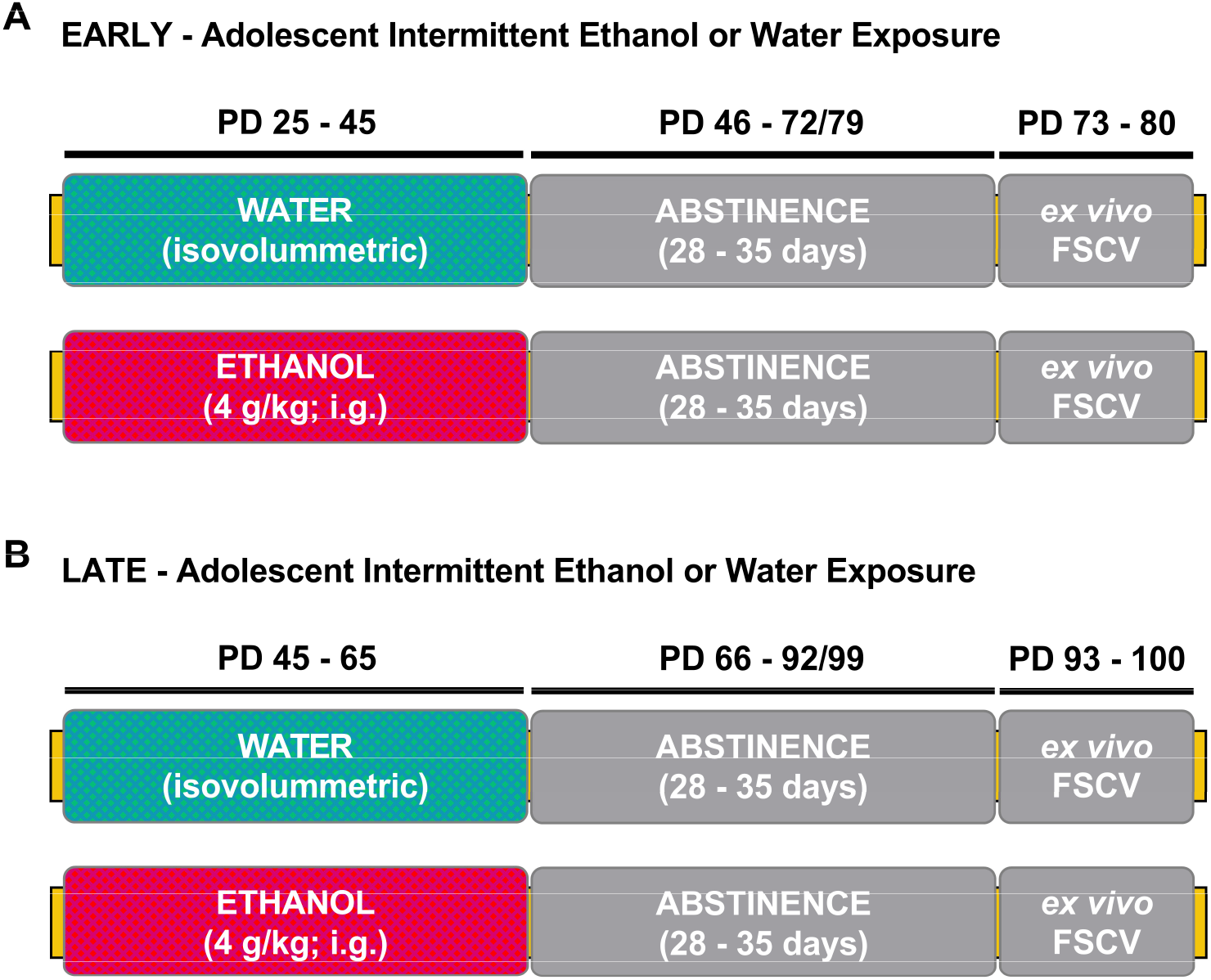
Timeline of experimental procedure. **(A)** Experimental manipulation during early adolescence. Male and female Sprague-Dawley rats were exposed intermittently, every other day for a total of 11 exposures, to water or ethanol (early AIE) via intragastric gavage between PD 25 and 45. After PD 45, all rats were subject to a period of protracted abstinence for 28-35 days, during which no manipulation took place. Ex vivo FSCV was conducted during adulthood, between PD 73 and 80. **(B)** Experimental manipulation during late adolescence. During PD 45-65, male and female rats were exposed intermittently, every other day for a total of 11 exposures, to either water or ethanol (late AIE) via intragastric gavage. After PD 65, all rats were subject to protracted abstinence for 28-35 days, during which no manipulation took place. Ex vivo FSCV was conducted during adulthood (between PD 93 – 100). PD, postnatal day; FSCV, Fast-Scan Cyclic Voltammetry.

### Ex vivo fast scan cyclic voltammetry

We used *ex vivo* FSCV to determine AIE-induced changes in dopamine kinetics and KOR function within the NAc core. All ex vivo FSCV recordings were conducted in adulthood after 28 – 35 days of forced abstinence, (PD 73 – 80 for early-AIE exposure and PD 93 – 100 for late-AIE exposure). *Ex vivo* FSCV procedures were similar to those used in previous studies [32]. Briefly, rats were anesthetized with isoflurane and euthanized via rapid decapitation. Brains were collected promptly and immersed in cold, oxygenated artificial cerebrospinal fluid (aCSF) and then sliced using a vibratome (VT1200 S, Leica BioSystems, Buffalo Grove, IL). Brain slices 400 μm thick containing the NAc core [38] were obtained and transferred to a recording chamber with a continuous flow of oxygenated aCSF (32°C). Dopamine efflux was induced using a bipolar stimulating electrode (8IMS3033SPCE, Plastics One, Roanoke, VA, USA) and measured using a carbon fiber recording electrode (~150 μm length, 7 μM radius; C 005722/6, Goodfellow, UK). The recording and stimulating electrodes were placed approximately 100 μm apart. The stimulating electrode was placed on the slice’s surface while the recording electrode was placed approximately 100 μm below the surface of the slice. Dopamine efflux was induced through the use of a single, rectangular, 4.0-ms duration electrical pulse (350 μA, monophasic, inter-stimulus interval: 300 seconds). To detect dopamine release, a triangular waveform (−0.4 to +1.2 to −0.4V vs. silver/silver chloride, 400V/sec) was applied every 100 ms to the recording electrode. Baseline dopamine recordings were taken prior to the bath application of any drug. Once stable baseline recordings were obtained, the KOR agonist U50,488 was bath applied in cumulative concentrations (0, 0.01, 0.03, 0.1, 0.3, 1.0 μM) to determine AIE-associated changes to KOR-mediated dopamine release. Recording electrodes used for each experiment were calibrated using a 3.0 μM concentration of dopamine in order to quantify the measured dopamine release and dopamine kinetics. FSCV recordings were later analyzed using Demon Voltammetry and Analysis software [39]. Dopamine kinetics were determined through modeling the stabilized signals using Michaelis-Menten kinetics [40].

### Statistical analysis

All data in this study were analyzed using GraphPad Prism 8 (GraphPad Software, La Jolla, CA, USA). Because our main focus was on examining the impact of AIE exposure on dopamine kinetics in males and females, and because the animals exposed in early and late adolescence were tested at different ages in adulthood, we analyzed the data separately for the early and late exposure groups. For each age of exposure, the dependent variables dopamine release and reuptake rate were analyzed using separate analysis for each measure using a 2 (adolescent exposure: water, AIE) X 2 (sex) analyses of variance (ANOVAs). Percent baseline dopamine release in response to a selective kappa agonist U50,488 was analyzed separately for each sex using a 2 (adolescent exposure: water, AIE) X 5 (U50,488 concentration: 0.01, 0.03, 0.1, 1.0, 3.0 μM) repeated measures ANOVA, with U50,488 concentration treated as a repeated measure. All main effects and interactions were explored using Holm-Sidak’s pairwise *post-hoc* analyses. Non-linear regression analysis was used to calculate EC50 in order to examine shifts in potency. Maximal change in dopamine release was calculated to examine changes in efficacy. Student’s *t*-tests were used to determine AIE exposure-induced changes in potency and efficacy within each age of exposure/sex condition. All data are reported as mean ± standard error of the mean. The significance level for all statistical measures was set at *p* < 0.05.

## RESULTS

### Late-AIE exposure disrupts dopamine release in male and female rats

In order to identify the impact of AIE exposure on dopamine kinetics in the NAc core, we measured electrically stimulated dopamine release and reuptake rate in adult male and female rats exposed to AIE or water during early or late adolescence using *ex vivo* FSCV. We assessed the early exposure groups separately (male: water = 6, AIE = 7; female: water = 7, AIE = 8) and late (male: water = 6, AIE = 6; female: water = 6, AIE = 8). **Figure 2A** shows representative traces of transient accumbal dopamine signals in response to a single pulse electrical stimulation in adult male (left) and adult female (right) rats exposed to water or ethanol during early adolescence. Baseline dopamine release (**Figure 2B**) and rate of uptake (**Figure 2C**) did not differ as a function of early adolescent exposure in adult males and females.

**Figure 2:**
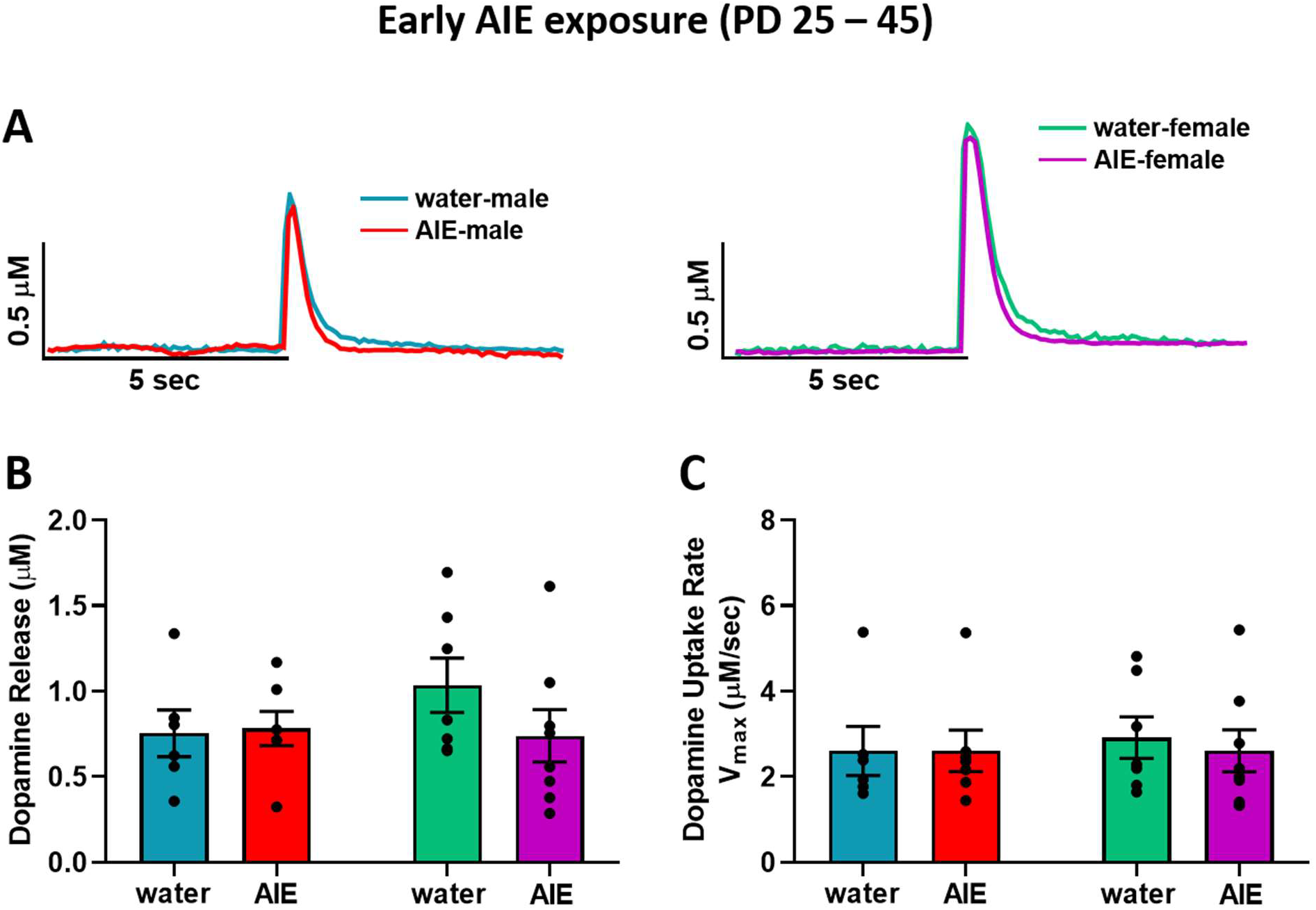
Baseline dopamine kinetics in the NAc core after early adolescent exposure (PD 25 – 45). **(A)** Representative transient dopamine signals in response to a single pulse electrical stimulation indicating concentration of dopamine released over time from the NAc core of one rat in each early exposure group (left: water-exposed male, blue; AIE-exposed male, red | right: water-exposed female, green; AIE-exposed female, purple). **(B)** Early AIE exposure had no effect on evoked dopamine release in male and female rats. **(C)** Similarly, the uptake rate of dopamine was not altered following early AIE exposure in both males and females. NAc, nucleus accumbens; AIE, adolescent intermittent ethanol.

**Figure 3A** shows representative traces of evoked dopamine response in the NAc core of adult male (left) and female (right) rats exposed to water and AIE during late adolescence. In contrast to early adolescent exposure, AIE exposure during late adolescence resulted in altered dopamine release (**Figure 3B**). A two-way ANOVA of dopamine release revealed an interaction between late adolescent exposure (water vs. AIE) and sex (*F*_(1,22)_=10.5; *p* < 0.01), with late AIE-exposure resulting in significantly lower dopamine release relative to water-exposed controls in females, while leading to elevated dopamine release in males in comparison to water-exposed controls (*p* < 0.05). In addition, water-exposed females showed significantly greater dopamine release compared to water-exposed males (*p* < 0.05). Although the ANOVA of dopamine uptake rate revealed a significant exposure by sex interaction (**Figure 3C**; *F*_(1,22)_ = 5.85; *p* < 0.05), *post-hoc* analyses did not identify any significant differences between water and AIE-exposed males and females. Collectively, the effects of AIE on dopamine release in NAc core appear to be impacted by exposure timing and sex.

**Figure 3:**
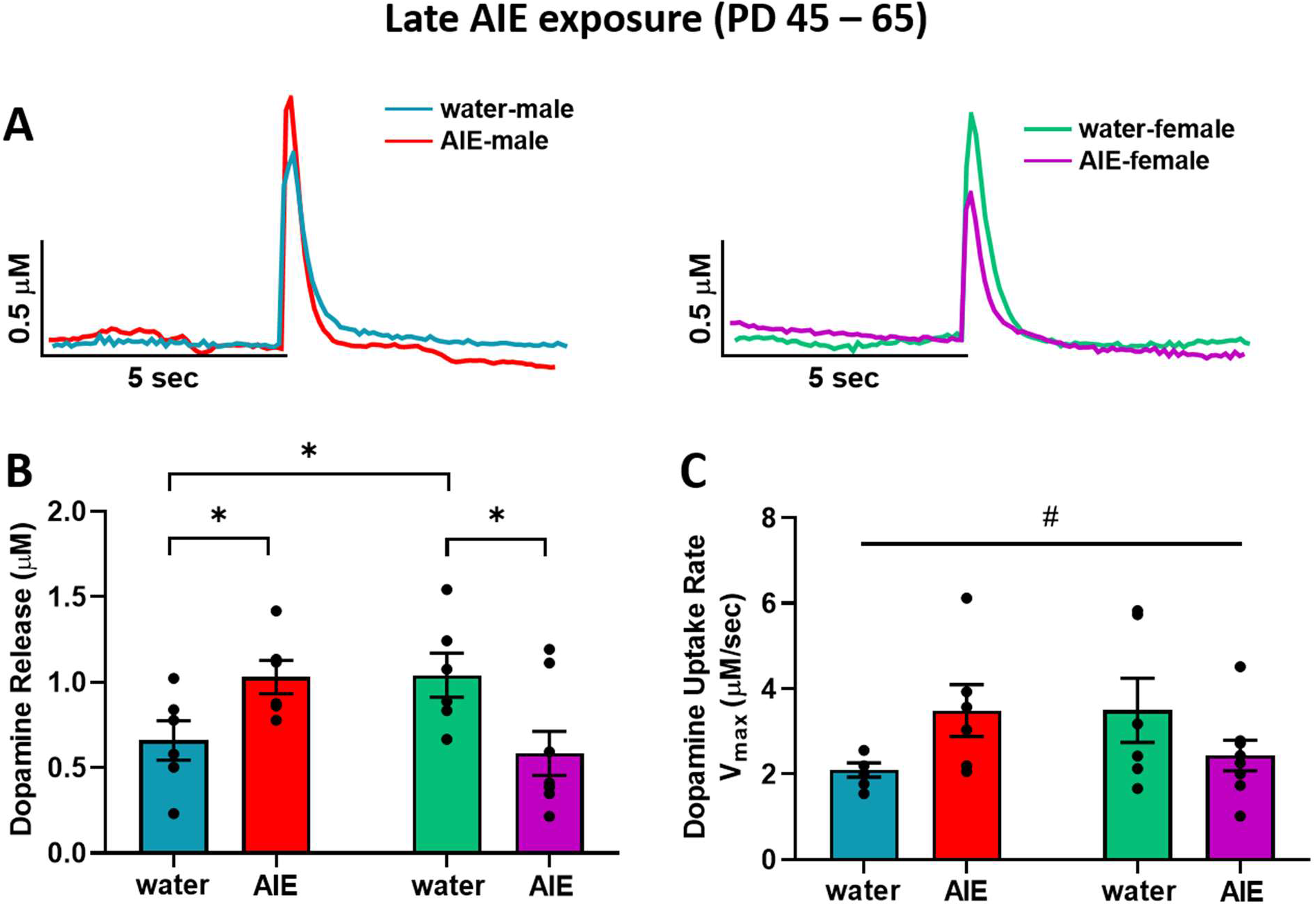
Baseline dopamine kinetics in the NAc core after late adolescent exposure (PD 45 – 65). **(A)** Representative transient dopamine signals in response to a single pulse electrical stimulation indicating concentration of dopamine released over time from the NAc core of one rat in each late exposure group (left: water-exposed male, blue; AIE-exposed male, red | right: water-exposed female, green; AIE-exposed female, purple). **(B)** Dopamine release in water-exposed females was significantly greater than dopamine release in water-exposed males. In males, late AIE-exposure (red bar) led to greater stimulated dopamine release compared to water exposed controls (blue bar). In females, late AIE-exposed rats (purple bar) exhibited lower stimulated dopamine release in comparison to water-exposed controls (green). **(C)** Analysis of the impact of AIE exposure on dopamine uptake rate revealed an interaction between sex and exposure. NAc, nucleus accumbens; AIE, adolescent intermittent ethanol. * - a significant difference between water and AIE-exposed rats within sex or between sexes in water-exposed groups where p<0.05; # - sex by treatment interaction where p<0.05.

### AIE exposure affects accumbal KOR function in an age- and sex-dependent manner

In order to determine the effects of AIE exposure on KOR function in the NAc core, we measured the functional responsiveness of KORs in water- and AIE-exposed male and female rats by measuring dopamine release following bath application of cumulative concentrations of a selective KOR agonist, U50,488 (0.01 – 1.0 μM), to accumbal slices. To further assess if AIE exposure affected potency and efficacy, we calculated the EC_50_ of U50,488 and the maximum effect of KOR activation on dopamine inhibition, respectively. In early exposure males (**Figure 4A**), U50,488 significantly decreased dopamine release as evidenced by a main effect of U50,488 concentration (F_(4,44)_ = 142.8; p < 0.0001), with no significant difference between early AIE-exposed and control groups. Likewise, no differences between early AIE-exposed and water-exposed males were observed in the EC50 (**Figure 4B**) and maximum effect (**Figure 4C**) of U50,488. In contrast, AIE exposure during early adolescence augmented KOR function in female rats relative to the water-exposed control group (**Figure 4D**). The ANOVA revealed main effects of early adolescent exposure (water vs. AIE; *F*_(1,13)_ = 10.4; *p* < 0.01) and U50,488 concentration (*F*_(4,52)_ = 157.3; *p* < 0.0001). U50,488 dose-dependently decreased dopamine release, and early AIE-exposed females demonstrated more pronounced decreases than their water-exposed counterparts, especially at 0.03, 0.10, 0.30, and 1.0 μM of U50,488 (*p* < 0.05), suggesting augmented KOR function in response to the agonist mediation activation. This hyperfunction of KORs was driven by an increase in potency (**Figure 4E**; *t*_13_ = 3.14; *p* < 0.01) and efficacy (**Figure 4F**; *t*_13_ = 2.55; *p* < 0.05). These data suggest that the consequences of AIE exposure during early adolescence followed by protracted abstinence differ between males and females; specifically, AIE exposure increases KOR function selectively in female rats.

**Figure 4:**
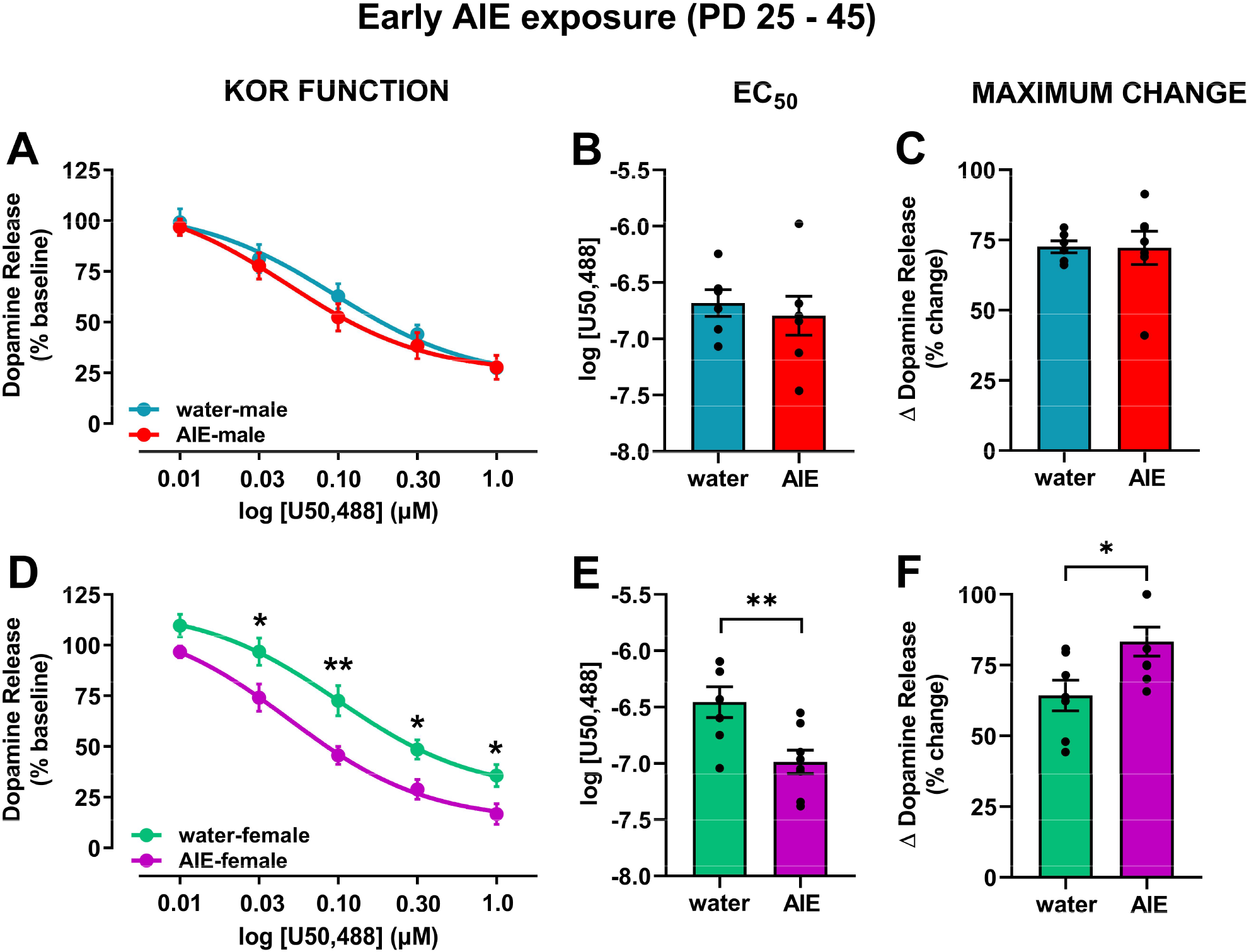
Impact of early AIE-exposure on KOR function and its regulation of dopamine release. The KOR agonist U50,488 was bath-applied to the tissue in cumulative concentrations and electrically stimulated dopamine release was recorded at each concentration in male (A – C) and female (D – E) rats. **(A)** In males, cumulative concentrations of U50,488 decreased dopamine release regardless of adolescent exposure to water or ethanol. **(B)** No differences in the EC50 for U50,488 were evident between water- and AIE-exposed exposed males. **(C)** Similarly, no differences in maximal change in dopamine release were found between water- and AIE-exposed male rats. **(D)** In females, both water- and AIE-exposed animals showed KOR-activation mediated inhibition of dopamine release; however, this effect was enhanced in AIE-relative to water-exposed rats at 0.03, 0.10, 0.30, and 1.0 μM of U50,488. These data suggest that early AIE-exposure resulted in augmented KOR function. Early AIE exposure-associated potentiation of KOR function was driven by increased potency, as shown by reduced EC50 of U50,488 **(E)**, as well as increased efficacy as shown by significantly greater maximum change in dopamine release **(F)** in early AIE-exposed females compared to water-exposed controls. Significant difference between water and AIE-exposed rats within sex: *p<0.05; **p<0.01. KOR, kappa opioid receptor; NAc, nucleus accumbens; AIE, adolescent intermittent ethanol; EC50, effective concentration of drug causing a half-maximal effect.

Contrary to the early-AIE exposure, late-AIE exposure followed by protracted abstinence attenuated KOR function in male rats with no change observed in females (Figure 5). In males, the ANOVA comparing KOR-activation mediated inhibition of dopamine release (**Figure 5A**) revealed main effects of exposure (water vs. AIE; *F*_(1,10)_ = 10.8; *p* < 0.01) and U50,488 concentration (*F*_(4,40)_ = 63.7; *p* < 0.0001). As expected, U50,488 concentration-dependently decreased dopamine release in both groups; however, late AIE-exposed males showed less pronounced decreases than their water-exposed counterparts, suggesting reduced functional response of the KORs to the U50,488 mediated activation in late-AIE males. *Post-hoc* comparisons showed a significant difference at the 0.03 μM and 0.10 μM concentrations of U50,488 (*p* < 0.01). This hypofunction of KORs was driven by a shift in potency (**Figure 5B**; *t*_10_=3.00; *p* < 0.05), but no difference in efficacy (**Figure 5C**; t_10_=1.35; p = 0.21). In females, the ANOVA comparing KOR-mediated inhibition of dopamine release showed a main effects of U50,488 concentration (**Figure 5D**; *F*_(4,48)_ = 107.5; *p* < 0.0001). However, AIE exposure during late adolescence in female rats did not result in altered KOR function in adulthood. As expected, no difference in efficacy and potency were observed between water- and AIE-exposed rats (**Figures 5E and F**).

**Figure 5:**
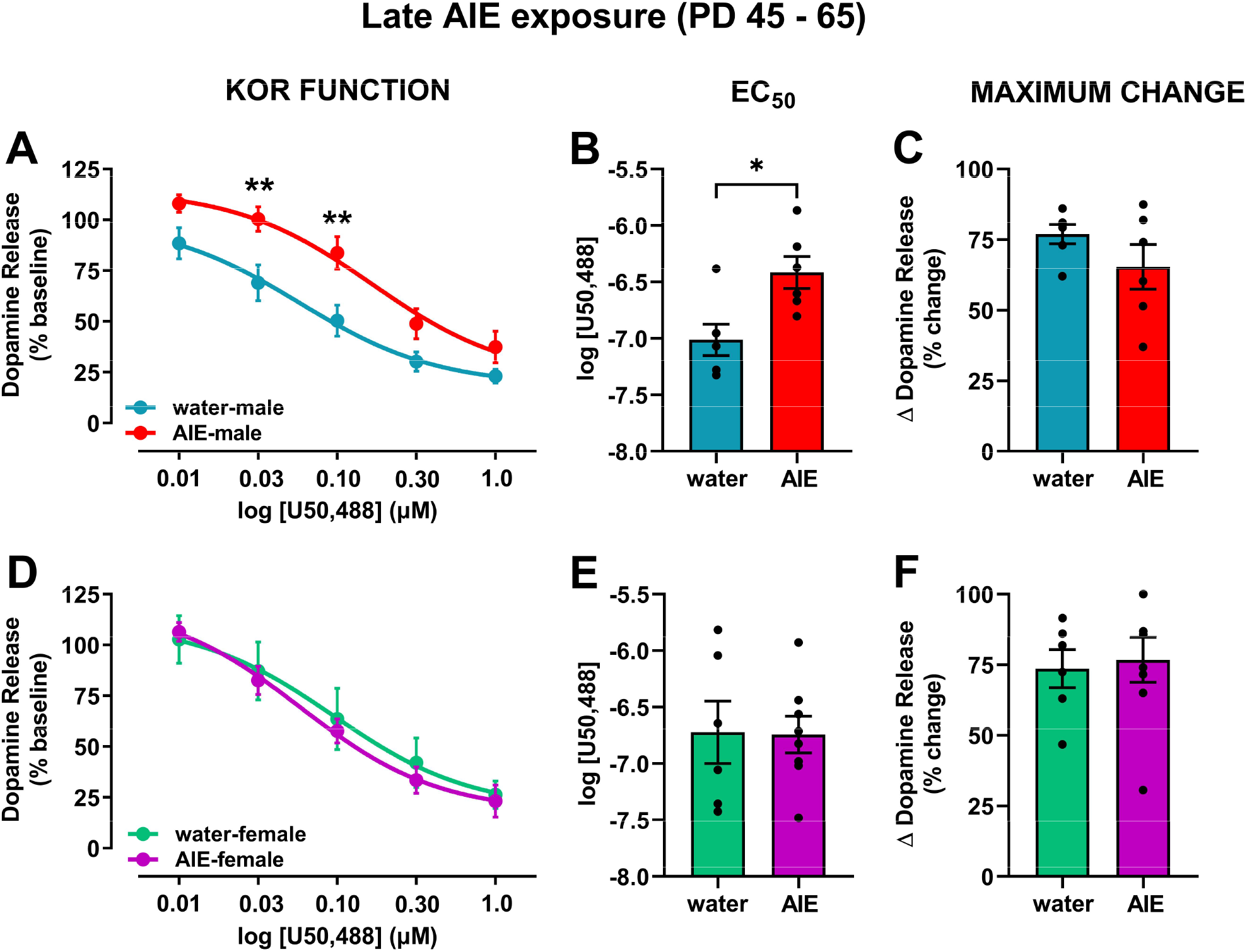
Impact of late AIE-exposure on KOR function and its regulation of dopamine release. The KOR agonist U50,488 was bath-applied to the tissue in cumulative concentrations and electrically stimulated dopamine release was recorded at each concentration in male (A – C) and female (D – E) rats. **(A)** In males, U50,488 concentration-dependently inhibited stimulated dopamine release in both AIE and water exposed rats; however, this effect was significantly attenuated in rats exposed to AIE, particularly at 0.03 and 0.1 μM of U50,488. These data suggest that late AIE-exposure reduced KOR function in male rats. **(B)** The AIE-associated attenuation in KOR function was driven by reduced potency as shown by augmented EC50 of U50,488 in late AIE-exposed male rats compared to water-exposed controls. **(C)** No differences were observed in the maximal effect of U50,488 on dopamine release between the AIE- and water-exposed groups, indicating that efficacy did not drive the changes in KOR function. **(D)** In females, bath application of U50,488 decreased stimulated DA release in a similar manner for both water- and AIE-exposed rats. No differences in potency **(E)** and efficacy **(F)** were observed across the two groups. Significant difference between water and AIE-exposed rats within sex: *p<0.05; **p<0.01; KOR, kappa opioid receptor; NAc, nucleus accumbens; AIE, adolescent intermittent ethanol; EC50, effective concentration of drug causing a half-maximal effect.

## DISCUSSION

In the current study, our data showed that AIE exposure-induced changes in dopamine transmission and KOR function were dependent on age of exposure as well as sex. We observed a sexually dimorphic effect of late AIE exposure, on baseline dopamine release: while stimulated dopamine release was greater in AIE-exposed males, it was lower in AIE-exposed females when compared to their respective control counterparts. These differences were not observed in early-exposure rats. In addition, neither early nor late AIE exposure influenced rate of dopamine uptake in the NAc. With regard to KOR function, our data showed a reduction in KOR function in late AIE-exposed males and an augmentation of KOR function in early AIE-exposed females, when compared to their respective water-exposed controls. We did not observe any changes in KOR function in late AIE-exposed females or early AIE-exposed males. Overall, these data suggest that the long-lasting impact of AIE on dopamine transmission and KOR function is dependent upon age of exposure and sex.

Adolescence is a critical period for maturation of the dopamine system [17]. Dopamine neurons in the VTA exhibit a greater firing rate in adolescent compared to adult rats [16,17]. Dopamine neuron firing increases from birth to early- and mid-adolescence (PD 42 – 48, period of peak dopamine firing rate), followed by a decline in activity until adulthood (> PD 70); forming an inverted U-curve in male rats [17,41–45]. The greater activity in adolescence seems to be driven by elevated tyrosine hydroxylase (TH) levels in midbrain tissue [16]. It is possible that in male rats, ethanol exposure during late adolescence interferes with the typical pruning process resulting in a maintenance of elevated adolescent-like dopamine neuron activity, as shown by greater stimulated dopamine release in late AIE-exposed versus water-exposed rats in the current study. The interference in the pruning process may be a result of intermittent ethanol-mediated excitation of these developing neurons during every ethanol exposure, i.e., the excitation surges caused by each ethanol exposure may result in potentiated dopamine release at baseline. Data in the current study showing increased dopamine release in late-AIE exposed males are congruent with a previous study showing elevated levels of dopamine in male rats exposed to ethanol from PD 30 to 50 [25]. These changes are likely not modulated by dopamine uptake as there are no AIE-induced persistent changes in the rate of dopamine uptake in adulthood [25]. In addition, it is important to note that the effects of ethanol on dopamine regulation primarily occur through its actions on release and not uptake [46]. Because the dopamine system is in a pre-pruning/growing state during early-mid adolescence, the ethanol-mediated excitation of dopamine neurons likely has minimal maladaptive changes in the dopamine system further leading to ‘normal’ baseline release in adulthood. Notably, data from the current study are congruent with a study that implemented *in vivo* FSCV and likewise found no change in baseline dopamine release in early AIE-exposed male rats following a 25-day forced abstinence period [26].

The development of the dopamine system has been studied relatively less in females. Developmental studies discussed in this paragraph were conducted in females only. Expression of Nurr1, a genetic transcription factor that activates transcription of dopamine and the vesicular monoamine transporters, increases between PD 28 and 60 in female rats [47]. Furthermore, in adulthood, Nurr1 is crucial in maintenance and survival of dopamine neurons [48,49]. On the other hand, expression of Pitx3, a transcription factor observed in mesolimbic dopamine neurons and involved in activating transcription of TH was observed to steadily decline with age (PD 7 – 60) [47]. Similarly, TH immunoreactivity also showed an age-related decline throughout early adolescence and into young adulthood [47]. These data suggest that maturation and pruning of the dopamine system likely occurs much earlier in females compared to maturation in males as described earlier. Therefore, it is possible that the late adolescent female neurochemistry is comparable to adult male neurochemistry resulting in similar chronic ethanol exposure-mediated changes to the dopamine system. Chronic intermittent ethanol-exposed male mice exhibit lower stimulated dopamine release at baseline compared to controls, up to 72 hours post cessation of the last exposure [21,22]. These data may explain the reduced dopamine levels in late AIE-exposed females in the current study. Because early AIE exposure ends during the developmental phase, the system may compensate and recover ‘normal’ dopamine release. However, late AIE exposure ends towards the end of the developmental phase, likely arresting the dopamine system in an altered state. Future experiments should look at the impact of these AIE exposures during acute withdrawal to test these hypotheses.

As mentioned earlier, KORs on dopamine neuron terminals have inhibitory control over dopamine release. Many studies examining the impact of chronic ethanol exposure in adult animals show ethanol-exposure mediated augmentation in KOR function. Specifically, studies demonstrate greater KOR-mediated inhibition of dopamine release in CIE-exposed adult male mice [21,22]. Moreover, inhibition of KORs attenuates ethanol consumption selectively in dependent animals [21,50]. This interaction between KORs and dopamine prompted us to determine whether the baseline changes we observed in late AIE-exposed rats were driven by AIE-associated changes in KOR function and their control over dopamine release. Interestingly, KOR-mediated inhibition of dopamine release was significantly lower in late AIE-exposed males relative to water-exposed controls, suggesting an attenuation in KOR function. This attenuation potentially explains the greater stimulated dopamine in late AIE-exposed exposed males relative to their water-exposed counterparts. Conversely, we did not find a significant difference in KOR function between late AIE- and water-exposed female rats, suggesting that the lower dopamine release observed in late AIE-exposed females is likely driven by a mechanism not dependent on KORs. Nevertheless, our data show that early AIE exposure in females resulted in augmented KOR function as shown by greater inhibition of dopamine release when compared to water-exposed controls. This effect was not observed in early AIE-exposed males.

Ontogenetically, mRNA for KORs has been observed as early as gestational day (G) 12 – PD 9, and the system seems to be fully matured by PD 21 – 25 [51]. Most studies show that the KOR mRNA expression and binding increases or fluctuates up to PD 9 – 10 followed by stable levels until PD 22 – 40 [52,53]. One study observed KOR mRNA expression levels to be significantly lower at PD 5 compared to adulthood [54]. It is important to note however, that the inhibitory effect of KORs on dopamine release in the NAc reduced from PD 7 to PD 21 [55]. These studies using male rats suggest that the KOR system matures by early adolescence. The KOR system development in females is largely understudied, thus comparable information in females is unknown. It is possible that the sex and exposure age differences that we observe are likely driven by the interaction between KORs and their ability to inhibit dopamine release and that AIE exposure at distinct time points (early vs. late adolescence) differentially affects KOR control over dopamine release. More research in this area is necessary to fully understand the underlying mechanisms.

Ethanol exposure during various developmental stages also differentially impacts KOR mRNA and function. Studies have shown that repeated prenatal ethanol exposure between G 17 – 20 resulted in higher levels of KOR mRNA in the NAc on PD 8 [56]. In contrast, similar prenatal ethanol exposure decreased KOR protein levels in the NAc on PD 14-15 [57]. Studies in neonatal rats have shown that activation of KORs is necessary to produce behavioral reinforcing effects of ethanol [58,59]. In addition, KOR inhibition blocked operant responding to ethanol in preweanling rats assessed on PD 14 – 17 [60]. Together these studies suggest that the relationship between ethanol and KORs is bidirectional; that is, ethanol exposure affects KOR function, KOR mRNA, and protein expression. In turn, KOR function contributes to behavioral effects of ethanol. These effects, however, are age- and sex-dependent. While inhibition of KORs in non-dependent adult male rats promoted ethanol intake, no changes were observed in adolescent (PD 25 – 38) rats. In contrast, KOR-inhibition reduced ethanol intake in adult females, with no changes evident in their adolescent counterparts [35]. Although these data suggest that KORs likely do not play a major role in adolescent ethanol consumption, it is important to note that ethanol exposure used in this study was mild and KOR inhibition effects on drinking were examined in non-dependent rats. Repeated exposure to ethanol, as used in the prenatal experiments discussed earlier, may be necessary to induce long-term changes in KOR function across the two sexes.

While prenatal ethanol exposure did not result in sex differences [56,57], it is possible that ethanol exposure during the peri-pubertal period may affect the dynorphin/KOR system in adulthood. Secretion of dynorphin in the hypothalamus reduces at the onset of puberty in female rats [61]. Alcohol exposure between PD 27 and 33 in female rats, however, resulted in elevated synthesis of dynorphin in this region [62]. Thus, it is possible that exposure to ethanol around the onset of puberty in females (PD 30 – 32) [63] results in retention of adolescent-typical enhancement in KOR function evident in the early AIE exposure female cohort in the current study. The KOR system is likely arrested in this altered state as a result of excessive ethanol exposure, even after termination of ethanol administration. Because late-AIE exposure occurred after puberty in the late-AIE exposed females, this effect was not observed in this group. It is possible that because early-AIE exposure in males ends right around the onset of puberty (PD 40 – 44) [63], with very few exposures occurring during this time, dynorphin levels might not be influenced, resulting in typical KOR function in adulthood. The augmented KOR function observed following late-AIE exposure in males is likely not related to puberty as most exposures occurred after sexual maturation. Further studies in adolescent males and females are needed for investigation of adolescent-typical KOR function pre- and during pubertal maturation, given that the role of puberty in KOR function is still not well documented and understood.

KOR function can also be altered as a result of changes in efficacy or potency. Our data showed that the augmented KOR function in early AIE-exposed females was a result of augmented efficacy, as shown by greater maximum effect of KOR activation on inhibition of dopamine release, as well as potency, as shown by lower EC50 of U50,488. On the other hand, the attenuated KOR function observed in late AIE-exposed males was driven by a reduction in potency of the receptor. These data suggest that these KOR changes in males and females are mechanistically distinct.

In summary, the results of the current study showed that while early AIE exposure followed by protracted abstinence did not affect dopamine kinetics in male or female rats, KOR-activation mediated inhibition of dopamine release was observed to be augmented in early AIE-exposed females only – an effect driven by enhanced potency and efficacy of the receptor. On the contrary, late AIE exposure followed by protracted abstinence resulted in facilitation of stimulated dopamine release in males and attenuation in females at baseline. Moreover, we observed a reduction in KOR-activation mediated inhibition of dopamine release, driven by enhanced potency selectively in males. Though we did not test direct mechanisms in this study, it is possible that the facilitation in stimulated dopamine release was driven by reduced KOR function in late AIE-exposed male rats. This relationship was not observed in early or late AIE-exposed females. Together, these data suggest that the impact of AIE on dopamine transmission and KOR function is exposure age and sex dependent, suggesting that AIE-associated changes in males and females may be driven by distinct mechanisms. Further research is necessary to elucidate the exact mechanisms that drive these neural adaptations.

## AUTHOR CONTRIBUTIONS

**Conceptualization**, Anushree N Karkhanis; **Data curation**, Mary B Spodnick, Raymond T Amirault and Trevor T Towner; **Formal analysis**, Mary B Spodnick, Raymond T Amirault and Anushree N Karkhanis; **Funding acquisition**, Elena I Varlinskaya, Linda P Spear and Anushree N Karkhanis; **Investigation**, Mary B Spodnick, Raymond T Amirault, Trevor T Towner, Elena I Varlinskaya, Linda P Spear and Anushree N Karkhanis; **Project administration**, Anushree N Karkhanis; **Resources**, Elena I Varlinskaya, Linda P Spear and Anushree N Karkhanis; **Supervision**, Anushree N Karkhanis; **Validation**, Anushree N Karkhanis; **Visualization**, Elena I Varlinskaya, Linda P Spear and Anushree N Karkhanis; **Writing – original draft**, Anushree N Karkhanis; **Writing – review & editing**, Mary B Spodnick, Raymond T Amirault, Trevor T Towner, Elena I Varlinskaya, Linda P Spear and Anushree N Karkhanis.

## REFERENCES

1. Geels LM, Bartels M, van Beijsterveldt TCEM, Willemsen G, van der Aa N, Boomsma DI, et al. Trends in adolescent alcohol use: effects of age, sex and cohort on prevalence and heritability. Addiction. 2012;107:518–27.

2. Merikangas KR, McClair VL. Epidemiology of substance use disorders. Hum Genet. 2012;131:779–89.

3. Lipari RN. Key Substance Use and Mental Health Indicators in the United States: Results from the 2018 National Survey on Drug Use and Health. 2018;82.

4. Grant BF, Dawson DA. Age at onset of alcohol use and its association with DSM-IV alcohol abuse and dependence: results from the National Longitudinal Alcohol Epidemiologic Survey. J Subst Abuse. 1997;9:103–10.

5. Dawson DA, Goldstein RB, Chou SP, Ruan WJ, Grant BF. Age at First Drink and the First Incidence of Adult-Onset DSM-IV Alcohol Use Disorders. Alcoholism: Clinical and Experimental Research. 2008;32:2149–60.

6. Ehlers CL, Slutske WS, Gilder DA, Lau P, Wilhelmsen KC. Age at First Intoxication and Alcohol Use Disorders in Southwest California Indians. Alcoholism: Clinical and Experimental Research. 2006;30:1856–65.

7. DeWit DJ, Adlaf EM, Offord DR, Ogborne AC. Age at First Alcohol Use: A Risk Factor for the Development of Alcohol Disorders. AJP. American Psychiatric Publishing; 2000;157:745–50.

8. Kann L, McManus T, Harris WA, Shanklin SL, Flint KH, Queen B, et al. Youth Risk Behavior Surveillance - United States, 2017. MMWR Surveill Summ. 2018;67:1–114.

9. Adger H, Saha S. Alcohol Use Disorders in Adolescents. Pediatr Rev. 2013;34:103–14.

10. Spear LP. Effects of adolescent alcohol consumption on the brain and behaviour. Nature Reviews Neuroscience. Nature Publishing Group; 2018;19:197–214.

11. Van Skike CE, Diaz-Granados JL, Matthews DB. Chronic intermittent ethanol exposure produces persistent anxiety in adolescent and adult rats. Alcohol Clin Exp Res. 2015;39:262–71.

12. Varlinskaya EI, Hosová D, Towner T, Werner DF, Spear LP. Effects of chronic intermittent ethanol exposure during early and late adolescence on anxiety-like behaviors and behavioral flexibility in adulthood. Behav Brain Res. 2020;378:112292.

13. Varlinskaya EI, Truxell E, Spear LP. Chronic intermittent ethanol exposure during adolescence: effects on social behavior and ethanol sensitivity in adulthood. Alcohol. 2014;48:433–44.

14. Gunaydin LA, Grosenick L, Finkelstein JC, Kauvar IV, Fenno LE, Adhikari A, et al. Natural neural projection dynamics underlying social behavior. Cell. 2014;157:1535–51.

15. Robinson DL, Heien MLAV, Wightman RM. Frequency of dopamine concentration transients increases in dorsal and ventral striatum of male rats during introduction of conspecifics. J Neurosci. 2002;22:10477–86.

16. McCutcheon JE, Conrad KL, Carr SB, Ford KA, McGehee DS, Marinelli M. Dopamine neurons in the ventral tegmental area fire faster in adolescent rats than in adults. Journal of Neurophysiology. 2012;108:1620–30.

17. Marinelli M, McCutcheon JE. Heterogeneity of dopamine neuron activity across traits and states. Neuroscience. 2014;282:176–97.

18. Gessa GL, Muntoni F, Collu M, Vargiu L, Mereu G. Low doses of ethanol activate dopaminergic neurons in the ventral tegmental area. Brain Res. 1985;348:201–3.

19. Brodie MS, Shefner SA, Dunwiddie TV. Ethanol increases the firing rate of dopamine neurons of the rat ventral tegmental area in vitro. Brain Res. 1990;508:65–9.

20. Imperato A, Di Chiara G. Preferential stimulation of dopamine release in the nucleus accumbens of freely moving rats by ethanol. J Pharmacol Exp Ther. 1986;239:219–28.

21. Rose JH, Karkhanis AN, Chen R, Gioia D, Lopez MF, Becker HC, et al. Supersensitive Kappa Opioid Receptors Promotes Ethanol Withdrawal-Related Behaviors and Reduce Dopamine Signaling in the Nucleus Accumbens. Int J Neuropsychopharmacol. 2016;19.

22. Karkhanis AN, Huggins KN, Rose JH, Jones SR. Switch from excitatory to inhibitory actions of ethanol on dopamine levels after chronic exposure: Role of kappa opioid receptors. Neuropharmacology. 2016;110:190–7.

23. Diana M, Pistis M, Muntoni A, Gessa G. Mesolimbic dopaminergic reduction outlasts ethanol withdrawal syndrome: evidence of protracted abstinence. Neuroscience. 1996;71:411–5.

24. Carrara-Nascimento PF, Hoffmann LB, Flório JC, Planeta CS, Camarini R. Effects of Ethanol Exposure During Adolescence or Adulthood on Locomotor Sensitization and Dopamine Levels in the Reward System. Front Behav Neurosci. 2020;14:31.

25. Badanich KA, Maldonado AM, Kirstein CL. Chronic Ethanol Exposure During Adolescence Increases Basal Dopamine in the Nucleus Accumbens Septi During Adulthood. Alcoholism: Clinical and Experimental Research. 2007;31:895–900.

26. Shnitko TA, Spear LP, Robinson DL. Adolescent binge-like alcohol alters sensitivity to acute alcohol effects on dopamine release in the nucleus accumbens of adult rats. Psychopharmacology (Berl). 2016;233:361–71.

27. Vetter-O’Hagen CS, Spear LP. The effects of gonadectomy on age- and sex-typical patterns of ethanol consumption in Sprague-Dawley rats. Alcohol Clin Exp Res. 2011;35:2039–49.

28. Spear LP. Adolescent alcohol exposure: Are there separable vulnerable periods within adolescence? Physiol Behav. 2015;148:122–30.

29. Sulzer D, Cragg SJ, Rice ME. Striatal dopamine neurotransmission: Regulation of release and uptake. Basal Ganglia. 2016;6:123–48.

30. Svingos AL, Chavkin C, Colago EE, Pickel VM. Major coexpression of kappa-opioid receptors and the dopamine transporter in nucleus accumbens axonal profiles. Synapse. 2001;42:185–92.

31. Di Chiara G, Imperato A. Opposite effects of mu and kappa opiate agonists on dopamine release in the nucleus accumbens and in the dorsal caudate of freely moving rats. J Pharmacol Exp Ther. 1988;244:1067–80.

32. Karkhanis AN, Rose JH, Weiner JL, Jones SR. Early-Life Social Isolation Stress Increases Kappa Opioid Receptor Responsiveness and Downregulates the Dopamine System. Neuropsychopharmacology. 2016;41:2263–74.

33. Spanagel R, Herz A, Shippenberg TS. Opposing tonically active endogenous opioid systems modulate the mesolimbic dopaminergic pathway. PNAS. National Academy of Sciences; 1992;89:2046–50.

34. Walker BM, Koob GF. Pharmacological evidence for a motivational role of kappa-opioid systems in ethanol dependence. Neuropsychopharmacology. 2008;33:643–52.

35. Morales M, Anderson RI, Spear LP, Varlinskaya EI. Effects of the kappa opioid receptor antagonist, nor-binaltorphimine, on ethanol intake: impact of age and sex. Dev Psychobiol. 2014;56:700–12.

36. Zorrilla EP. Multiparous species present problems (and possibilities) to developmentalists. Dev Psychobiol. 1997;30:141–50.

37. Spear LP. Timing Eclipses Amount: The Critical Importance of Intermittency in Alcohol Exposure Effects. Alcohol Clin Exp Res. 2020;44:806–13.

38. Paxinos G, Watson C. The Rat Brain in Stereotaxic Coordinates. Compact 6th Edition. New York, 400: Academic Press; 2005.

39. Yorgason JT, España RA, Jones SR. Demon voltammetry and analysis software: analysis of cocaine-induced alterations in dopamine signaling using multiple kinetic measures. J Neurosci Methods. 2011;202:158–64.

40. Ferris MJ, Calipari ES, Yorgason JT, Jones SR. Examining the complex regulation and drug-induced plasticity of dopamine release and uptake using voltammetry in brain slices. ACS Chem Neurosci. 2013;4:693–703.

41. Tepper JM, Trent F, Nakamura S. Postnatal development of the electrical activity of rat nigrostriatal dopaminergic neurons. Developmental Brain Research. 1990;54:21–33.

42. Pitts DK, Freeman AS, Chiodo LA. Dopamine neuron ontogeny: Electrophysiological Studies. Synapse. 1990;6:309–20.

43. McCutcheon JE, Marinelli M. Age matters. Eur J Neurosci. 2009;29:997–1014.

44. Philpot RM, Wecker L, Kirstein CL. Repeated ethanol exposure during adolescence alters the developmental trajectory of dopaminergic output from the nucleus accumbens septi. International Journal of Developmental Neuroscience. 2009;27:805–15.

45. Maldonado-Devincci AM, Badanich KA, Kirstein CL. Alcohol during adolescence selectively alters immediate and long-term behavior and neurochemistry. Alcohol. 2010;44:57–66.

46. Yim HJ, Gonzales RA. Ethanol-induced increases in dopamine extracellular concentration in rat nucleus accumbens are accounted for by increased release and not uptake inhibition. Alcohol. 2000;22:107–15.

47. Katunar MR, Saez T, Brusco A, Antonelli MC. Immunocytochemical expression of dopamine-related transcription factors Pitx3 and Nurr1 in prenatally stressed adult rats. J Neurosci Res. 2009;87:1014–22.

48. Prakash N, Wurst W. Genetic networks controlling the development of midbrain dopaminergic neurons. J Physiol. 2006;575:403–10.

49. Alavian KN, Scholz C, Simon HH. Transcriptional regulation of mesencephalic dopaminergic neurons: the full circle of life and death. Mov Disord. 2008;23:319–28.

50. Anderson RI, Lopez MF, Becker HC. Stress-Induced Enhancement of Ethanol Intake in C57BL/6J Mice with a History of Chronic Ethanol Exposure: Involvement of Kappa Opioid Receptors. Front Cell Neurosci [Internet]. Frontiers; 2016 [cited 2020 Jun 10];10. Available from: https://www.frontiersin.org/articles/10.3389/fncel.2016.00045/full

51. Georges F, Normand E, Bloch B, Le Moine C. Opioid receptor gene expression in the rat brain during ontogeny, with special reference to the mesostriatal system: an in situ hybridization study. Developmental Brain Research. 1998;109:187–99.

52. Winzer-Serhan UH, Chen Y, Leslie FM. Expression of opioid peptides and receptors in striatum and substantia nigra during rat brain development. J Chem Neuroanat. 2003;26:17–36.

53. Kornblum HI, Hurlbut DE, Leslie FM. Postnatal development of multiple opioid receptors in rat brain. Developmental Brain Research. 1987;37:21–41.

54. Stiene-Martin A, Knapp PE, Martin K, Gurwell JA, Ryan S, Thornton SR, et al. Opioid system diversity in developing neurons, astroglia, and oligodendroglia in the subventricular zone and striatum: Impact on gliogenesis in vivo. Glia. 2001;36:78–88.

55. De Vries TJ, Hogenboom F, Mulder AH, Schoffelmeer AN. Ontogeny of mu-, delta- and kappa-opioid receptors mediating inhibition of neurotransmitter release and adenylate cyclase activity in rat brain. Brain Res Dev Brain Res. 1990;54:63–9.

56. K B, T D. Endogenous Opioids as Substrates for Ethanol Intake in the Neonatal Rat: The Impact of Prenatal Ethanol Exposure on the Opioid Family in the Early Postnatal Period [Internet]. Physiology & behavior. Physiol Behav; 2015 [cited 2020 Jun 10]. Available from: https://pubmed.ncbi.nlm.nih.gov/25662024/?from_single_result=bordner+deak+2015

57. Nizhnikov ME, Pautassi RM, Carter JM, Landin JD, Varlinskaya EI, Bordner KA, et al. Brief Prenatal Ethanol Exposure Alters Behavioral Sensitivity to the Kappa Opioid Receptor Agonist (U62,066E) and Antagonist (Nor-BNI) and Reduces Kappa Opioid Receptor Expression. Alcoholism: Clinical and Experimental Research. 2014;38:1630–8.

58. Nizhnikov ME, Varlinskaya EI, Petrov ES, Spear NE. Reinforcing properties of ethanol in neonatal rats: involvement of the opioid system. Behav Neurosci. 2006;120:267–80.

59. Pautassi RM, Nizhnikov ME, Acevedo MB, Spear NE. Early role of the κ opioid receptor in ethanol-induced reinforcement. Physiol Behav. 2012;105:1231–41.

60. Miranda-Morales RS, Spear NE, Nizhnikov ME, Molina JC, Abate P. Role of mu, delta and kappa opioid receptors in ethanol-reinforced operant responding in infant rats. Behav Brain Res. 2012;234:267–77.

61. Dees WL, Hiney JK, Srivastava VK. Regulation of prepubertal dynorphin secretion in the medial basal hypothalamus of the female rat. J Neuroendocrinol. 2019;31:e12810.

62. Srivastava VK, Hiney JK, Dees WL. Alcohol Delays the Onset of Puberty in the Female Rat by Altering Key Hypothalamic Events. Alcohol Clin Exp Res. 2018;42:1166–76.

63. Vetter-O’Hagen CS, Spear LP. Hormonal and physical markers of puberty and their relationship to adolescent-typical novelty-directed behavior. Dev Psychobiol. 2012;54:523–35.

